# Rapid Whole Genome Sequencing Decreases Morbidity and Healthcare Cost of Hospitalized Infants

**DOI:** 10.1101/253534

**Authors:** Lauge Farnaes, Amber Hildreth, Nathaly M. Sweeney, Michelle M. Clark, Shimul Chowdhury, Shareef Nahas, Julie A. Cakici, Wendy Benson, Robert H. Kaplan, Richard Kronick, Matthew N. Bainbridge, Jennifer Friedman, Jeffrey J. Gold, Yan Ding, Narayanan Veeraraghavan, David Dimmock, Stephen F. kingsmore, on behalf of the RCIGM Investigators

## Abstract

**BACKGROUND:** Genetic disorders are a leading cause of morbidity and mortality in infants. Rapid Whole Genome Sequencing (rWGS) can diagnose genetic disorders in time to change acute medical or surgical management (clinical utility) and improve outcomes in acutely ill infants.

**METHODS:** Retrospective cohort study of acutely ill inpatient infants in a regional children’s hospital from July 2016–March 2017. Forty-two families received rWGS for etiologic diagnosis of genetic disorders. Probands received standard genetic testing as clinically indicated. Primary end-points were rate of diagnosis, clinical utility, and healthcare utilization. The latter was modelled in six infants by comparing actual utilization with matched historical controls and/or counterfactual utilization had rWGS been performed at different time points.

**FINDINGS:** The diagnostic sensitivity was 43% (eighteen of 42 infants) for rWGS and 10% (four of 42 infants) for standard of care (P=.0005). The rate of clinical utility for rWGS (31%, thirteen of 42 infants) was significantly greater than for standard of care (2%, one of 42; P=.0015). Eleven (26%) infants with diagnostic rWGS avoided morbidity, one had 43% reduction in likelihood of mortality, and one started palliative care. In six of the eleven infants, the changes in management reduced inpatient cost by $800, 000 to $2,000,000.

**DISCUSSION:** These findings replicate a prior study of the clinical utility of rWGS in acutely ill inpatient infants, and demonstrate improved outcomes and net healthcare savings. rWGS merits consideration as a first tier test in this setting.

## Introduction

Genetic disorders and congenital anomalies affect ~6% of live births, and are the leading reason for hospitalization and mortality in infants(1-4). Of 14% of US newborns admitted to neonatal intensive care units (NICU), those with genetic disorders have longer hospitalizations and higher resource utilization(1, 5). While early etiologic diagnosis in such infants enables optimal outcomes, it is exceptionally difficult to deliver for genetic diseases since they number over 8,000 and presentations are often atypical from classical descriptions(6). Moreover, they represent the leading cause of NICU and paediatric intensive care unit (PICU) mortality, with most deaths following palliative care decisions. Family counselling regarding palliative care often is impeded by absence of etiologic diagnosis(7).

Rapid whole genome sequencing (rWGS) provides a faster diagnosis, enabling precision medicine interventions in time to decrease the morbidity and mortality of infants with genetic diseases.(6) Furthermore, rWGS facilitates end-of-life care decisions that can alleviate suffering and aid the grieving process. However, published evidence demonstrating the effectiveness of rWGS in improving outcomes in infants is insufficient to endorse large-scale implementation(8); It is limited to case reports, and one retrospective study (Level III evidence)(9). Examination of reproducibility is imperative. Here, we report such an examination.

## Findings

Parents provided consent for 42 of 48 eligible infants (88%, figure 1). While the intent was trio rWGS (parents and affected infant), rWGS was performed on 29 trios and 1 quad (parents and two affected siblings), 9 mother-infant duos, and 3 singletons.

**Figure. 1.**
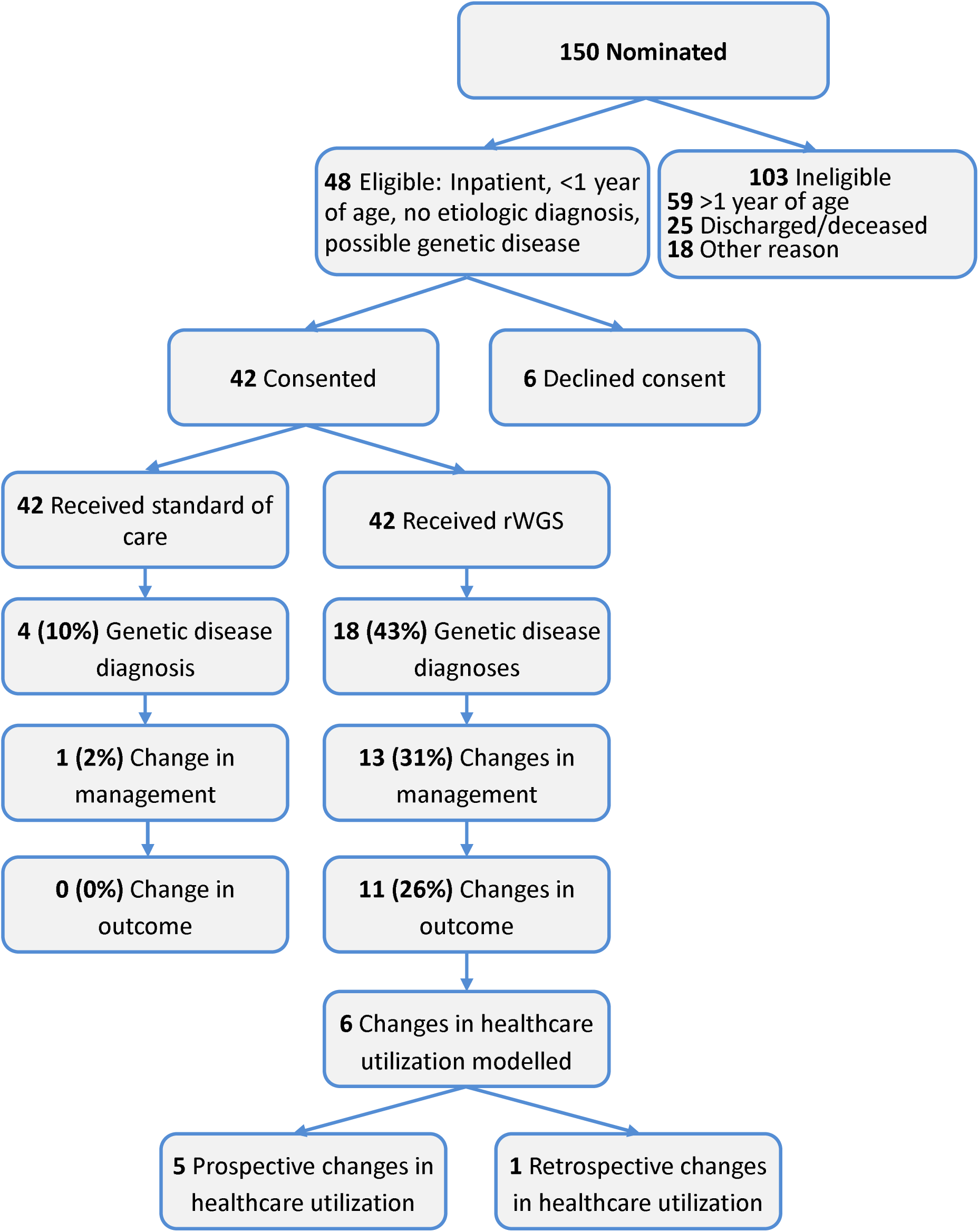
Flow diagram of the proportion of inpatient infants who were enrolled, received genetic disease diagnoses by rWGS or by standard tests, had consequent acute changes in management (precision medicine), resultant change in outcome, and analysis of impact on acute healthcare utilization.

Most probands were term births with normal weight (table 1). 59% were Hispanic/Latino. Consanguinity was uncommon (2%). The severity of illness was high: 71% were in a regional NICU, PICU or cardiovascular intensive care unit, 76% received respiratory support and 40% inotropic support. Disease phenotypes were highly diverse (tables 1, S1). Most presentations were complex, with several systems affected. The most common presentation was multiple congenital anomalies (29%).

**Table 1.**
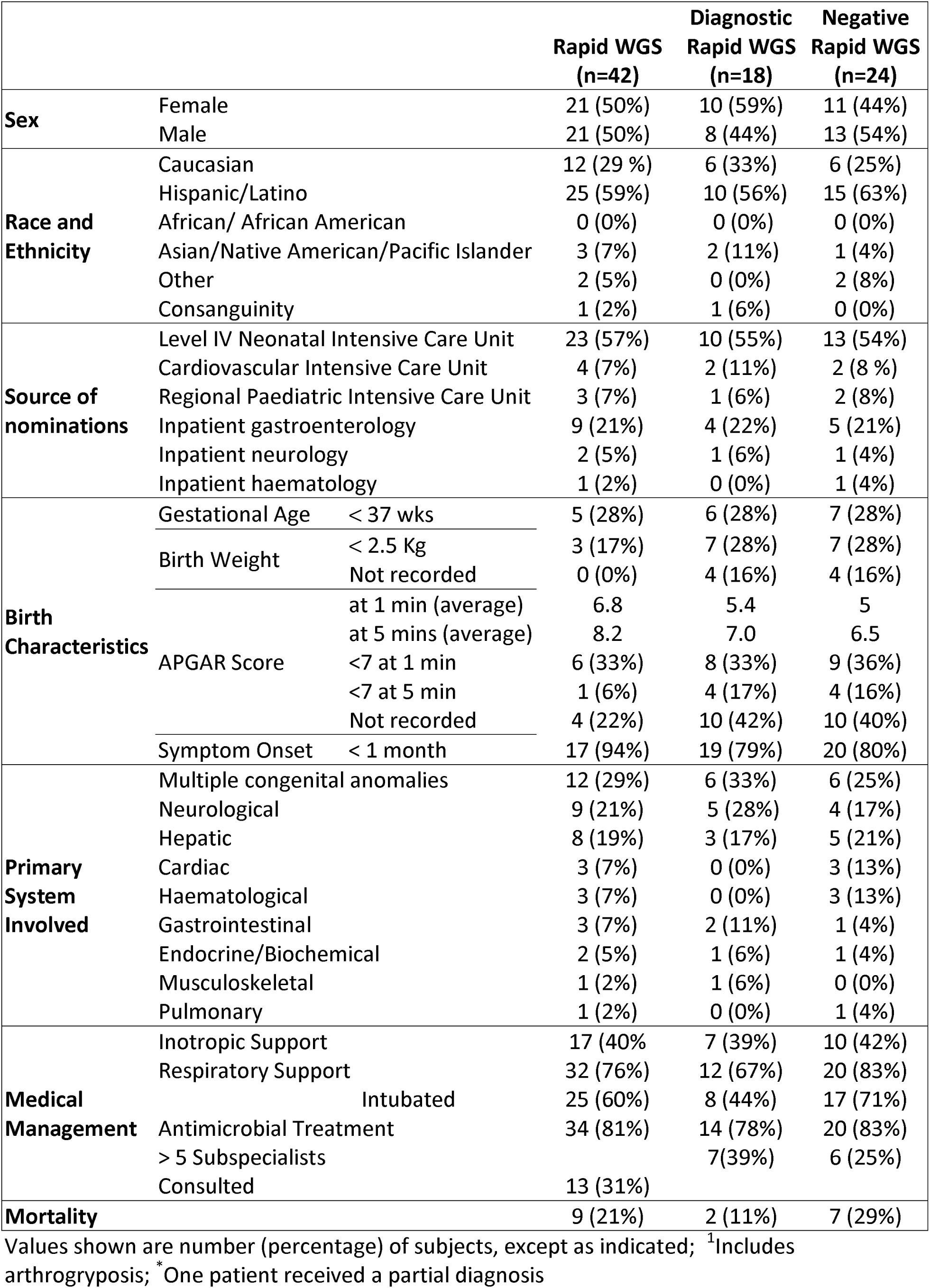
Demographic and clinical characteristics of the 42 proband, inpatient infants

While disease phenotypes were typically present at birth, the median age at nomination was 62 days (range 1– 301 days; Table S5). The median time from admission to nomination was 5 days. The range of the interval between admission and nomination varied from nomination at a prior admission (in two infants) to 194 days. The median enrolment interval was 5 days (range 0–62 days), and enrolment–receipt of proband blood sample was zero days (range 0-6 days).

## Diagnostic Sensitivity of rWGS and Standard of Care

rWGS diagnosed 19 genetic diseases in 18 of 42 infants (figure 1, table S3). All were confirmed with standard genetic tests. Concomitantly, 33 (79%) of 42 infants received 144 standard genetic tests, as clinically indicated (table S5). The most common was CMA (18 infants, 43%). The diagnostic sensitivity of standard of care (10%) was significantly less than rWGS (43%) (P=.0026). rWGS identified all diagnoses made by standard genetic testing. In addition, one infant was incidentally diagnosed with adenovirus infection by metagenomic analysis of WGS. Thus 19 (45%) infants received etiologic diagnoses by rWGS. No diagnoses were considered false positives by clinicians (diagnostic specificity 100%).

The most common presentations receiving diagnoses were multiple congenital anomalies (6 (50%) of 12 were diagnosed), and neurologic (5 (56%) of 9 were diagnosed; table 1). No demographic or clinical characteristic was significantly associated with diagnosis.

In 7 (17%) very ill infants, rWGS was performed with two day time to interpretation. The remaining 35 (83%) families received rWGS using less costly methods with five days to interpretation. In 6 families (33% of positive results) WGS established a diagnosis for which a specific treatment was available to prevent morbidity or mortality. Thus prompting results to be called out ahead of medically accepted confirmatory testing. All were subsequently confirmed, with specific therapy initiated before this confirmation.

## Clinical Utility

Specific changes in medical or surgical treatment occurred as a result of molecular diagnoses (clinical utility) in 13 (31%) of 42 infants receiving rWGS (including all who received provisional diagnoses, table 2). Conversely only one child (2%) received changes in care from standard genetic diagnostics (P=0.0015). Seven medications were started in five infants, and medications were discontinued in two (excluding discontinuation of empiric antibiotics prescribed in 82% of infants, Table 1). For example, infant 6056, was diagnosed with megacystic microcolon intestinal hypoperistalsis syndrome, which is lethal in most infants(15). Treatment with cisapride resulted in marked improvement in gut motility, such that intestine transplant may not be necessary(16). In four infants (22% of diagnosed), surgical procedures were changed. Three were major surgeries. *In toto*, rWGS-predicated precision medicine was judged to have prevented morbidity in eleven (61%) of eighteen diagnosed infants, compared with none by standard of care (table 2). Four infants (22% of diagnosed) avoided major morbidity (discussed below; Table 2). Mortality through median DOL 430 was two (11%) of eighteen diagnosed by rWGS (infant 6012 and 6034), compared with seven (29%) of 24 infants who did not receive diagnoses (table 1, S4). Mortality was bimodal, with four deaths at DOL 29-35, and five at DOL 209-561.

**Table 2.** Acute precision medicine interventions in thirteen of eighteen infants receiving genetic disease diagnoses and the resultant changes in outcomes.

| Infant ID | Causal Gene | Medication Change | Change in Surgery | Palliative Care Initiated | Imaging or Procedure Change | Morbidity Avoided | Mortality Avoided |
| --- | --- | --- | --- | --- | --- | --- | --- |
| 6011 | *NPC1* | Miglustat started |  |  |  | Neurologic damage delayed | - |
| 6012 | *ARID1B* |  |  | Yes |  | Further futile intensive care | - |
| 6014 | *NEB* |  | Avoided muscle Biopsy |  | Avoided EMG and NCS | Anaesthesia and muscle biopsy | - |
| 6018 | *POLR1C* |  |  |  | MRI of brain recommended |  | - |
| 6019 | *GABRA1* | Steroids weaned; confidence in therapy when readmitted |  |  | Avoided repeat EEG | Discontinuation of appropriate anti-epileptic at next admission | - |
| 6020 | *TPM1* |  | Cleared for cardiac transplant |  |  | Delay in heart transplant | - |
| 6021 | *PCDH19* | Ganaxolone started; Confidence in medications for child and sibling |  |  |  |  | - |
| 6024 | *PHEX* | Start phosphate and high-dose calcitriol |  |  |  | Development of rickets | - |
| 6026 | *JAG1* |  | Avoided Kasai hepatoportoenterostomy |  |  | Kasai and Liver Transplant | 83%-94% decrease |
| 6030 | *NF1* |  |  |  | Brain MRI and MR angiography of renal arteries | Potential early detection of *NF1* associated tumours | - |
| 6041 | *KCNQ2* | Carbamazepine started; phenobarbital weaned |  |  |  | Prolonged uncontrolled seizures with potential neurological damage | - |
| 6053 | *ABCC8* |  | Earlier partial pancreatectomy |  |  | 3 additional weeks of hypoglycaemia with potential neurological damage | - |
| 6056 | *ACTG2* | Started Cisapride |  |  |  |  |  |
| Total |  | 5 (28%) | 4 (22%) | 1 (6%) | 4 (22%) | 10 (56%) | 1 (6%) |

## Impact of rWGS-Associated Precision Medicine on Healthcare Utilization

Although appearing to reduce rather than increase costs, the impact of precision medicine could not be quantified in seven of thirteen infants without a group of matched historical controls and/or long-term follow up (see Supplementary Results). However, the short-term impact on healthcare utilization was quantified in the remaining six infants (table 3, Figure 1). Infant 6011 (previously published(11)) was admitted at seven weeks of age with persistent jaundice and poor weight gain. He was nominated for rWGS, but Spanish consent forms had not yet been approved. Despite extensive evaluation, the aetiology of cholestasis was not identified. He was discharged after eight days, but readmitted four days later due to α-fetoprotein levels of >200,000 ng/mL. rWGS revealed a diagnosis of Niemann-Pick disease, type C1 (NPC1). He started treatment with miglustat with plans for intrathecal cyclodextran and remains without neurologic symptoms(11). Using a modified Delphi method, the consensus of an international panel of paediatricians was that, had rWGS been performed on his first admission, the second admission would have been unnecessary.

**Table 3.** Effect of rWGS-based precision medicine on acute healthcare utilization in six infants and three matched controls. Change in Healthcare costs due to whole genome sequencing. Patient 6011 would have avoided second hospitalization if NPC1 diagnosis was made on first admission saving $27,004. Patient 6012 had compassionate withdrawal of care after diagnosis of ARID1B negating need for 42 day planned antibiotic course and continued intensive care stay with a projected cost of $327,506. Patient 6014 avoided a muscle biopsy and EMG and likely need for recovery in NICU from anaesthesia due to hypotonia saving $9,900. Patient 6026 avoided the need for a Kasai saving $44,451 and also avoided 43% increased likelihood of needing liver transplant if the Kasai had been performed resulting in net savings of $87,344 ($203,125 (cost of surgery and 90 days post-transplant) x 43%). Patient 6041 had diagnosis made 38 days earlier than case control who had standard work up one year earlier resulting in savings of $181,481. Patient 6053 had diagnosis of ABCC8 made 21 days earlier than what is the standard from the literature resulting in a net savings of $125,514 from shortened stay in NICU while trying to control severe hypoglycaemia. Total savings was $803,200 which was $128,555 less than actual cost of sequencing all 42 families.

| Subject ID | Presentation and modeled change in care | Gene | Time-to-Diagnosis, |  | Total Cost | Cost Avoided |
| --- | --- | --- | --- | --- | --- | --- |
|  |  |  | Days |  |  |  |
| 6011 | Cholestasis. 1<sup>st</sup> admision for etiologic Dx (8 days) | NPC1 | 7 | \$ | 25,278 | \$ 27,004 |
| | Cholestasis. 2<sup>nd</sup> admission for etiologic Dx (7 days) | | | \$ | 27,004 | |
| 6012 | Palliative care started DOL 250 | ARID1B | 26 | \$ | 1,949,438 | \$ 327,506 |
| | Palliative care started DOL 292 | | | \$ | 2,276,944 | |
| 6014 | Hypotonia. Avoided EMG, GA, muscle biopsy | NEB1 | 7 | \$ | 156,914 | \$ 9,900 |
| Ctrl 1 | EMG, GA, muscle biopsy |  |  |  |  |  |
| 6026 | Cholestasis and congenital heart disease. Avoided | JAG1 | 3 | \$ | 50,327 | \$ 131,795 |
| Ctrl 2 | Kasai hepatoportoenterostomy | | | \$ | 44,451 | |
| Avg Cost | Cost of Liver Transplant * 43% occurrence | | | \$ | 87,344 | |
| 6041 | Seizures. Diagnosis DOL 4 | KCNQ2 | 4 | \$ | 79,675 | \$ 181,481 |
| | Seizures. Diagnosis DOL 42 | | 42 | \$ | 261,156 | |
| 6053 | Hypoglycemia. Diagnosis DOL 12 | ABCC8 | 7 | \$ | 59,769 | \$ 125,514 |
| | Hypoglycemia. Diagnosis DOL 32 | | 28 | \$ | 185,283 | |
| Healthcare Savings | | | | | | \$ 803,200 |
| Cost of sequencing all 42 families | | | | | | \$ 674,645 |
| Net healthcare savings | | | | | | \$ 128,555 |

Ctrl: Control subject; Avg: Average.

Infant 6012 was admitted to the NICU at birth with congenital heart disease (Shone’s complex) and congenital diaphragmatic hernia. She had an extremely complicated course with multiple surgical interventions, including post-cardiac surgery extracorporeal membrane oxygenation, gastrostomy tube placement with Nissen fundoplication, tracheostomy for respiratory failure, and treatment for recurrent sepsis. She was enrolled on day of life (DOL) 224, shortly after the study commenced. A diagnosis of Coffin-Siris syndrome was reported on DOL 250, at which time she had septic shock, requiring inotropic support with multiple agents. A six week course of intravenous antibiotics had been planned. Upon diagnosis, in light of the prognosis, the family elected for comfort care, and the patient expired that day. Time to death after withdrawal of life-sustaining treatment is not commonly a protracted event(17, 18). The Delphi panel could not achieve consensus with respect to earlier election of palliative care if the diagnosis had been known soon after birth. However, there was consensus that without rWGS, NICU care and therapy for sepsis would have continued until resuscitation was no longer possible.

Neonate 6014 was admitted to the NICU at birth with respiratory distress. He was hypotonic and aspirated feeds, resulting in gastrostomy tube placement. He was enrolled on DOL 35, and a provisional diagnosis of Nemaline Myopathy was communicated on DOL 42. The panel consensus was that having the molecular diagnosis avoided the need for muscle biopsy, and that it was reasonable to compare projected costs of this procedure (from a matched control infant).

Infant 6026 was admitted to inpatient gastroenterology on DOL 76 with severe cholestasis. He was noted to have congenital heart disease. On DOL 78 he developed respiratory distress and metabolic acidosis, and was transferred to the PICU. He was scheduled for intraoperative cholangiogram with reflex to Kasai hepatoportoenterostomy on DOL 83 for evaluation of suspected biliary atresia (see supplementary results). He was enrolled on DOL 80 and received a provisional diagnosis of Alagille syndrome on DOL 83. This provisional diagnosis was reported to the operating room immediately before induction of general anaesthesia for intraoperative cholangiogram and Kasai procedure. The diagnosis was confirmed by CMA several days later. Control infant 2 was admitted to inpatient gastroenterology with cholestasis one month later, and underwent hepatoportoenterostomy for biliary atresia, costs were included for an expected additional two days of PICU recovery. Estimated rWGS-associated savings were $44,451. Given that 90% of infants with Alagille syndrome have abnormal intraoperative cholangiograms, the panel consensus was that he had 83% - 94% reduction in mortality as a result of cancellation of hepatoportoenterostomy, and 70% - 80% reduction in likelihood of liver transplant(19-21). Estimates of the cost of a subsequent liver transplant were based on an average charges of $812,500(22) given the typical 25% cost to charge ratio, estimated costs would be $203,125. Based on two previous publications(19, 20) which showed total of 14 out of 24 patients with Alagille who had a Kasai requiring liver transplant compared with 7 out of 46 patients with Alagille syndrome without Kasai requiring liver transplant it was estimated that Kasai increased rate of liver transplant by 43%. Consequently the likelihood weighted saving was $87,344 for liver transplant. Total estimated savings were $131,795.

Neonate 6041 was born by normal delivery at term, and immediately admitted to the NICU for tonic-myoclonic seizures. While seizures were largely controlled by phenobarbital, levetiracetam and topiramate, she was too sedated to tolerate oral feeds. Upon dose reduction, the seizures returned. Electroencephalogram revealed multifocal, epileptiform abnormalities and burst suppression. She was enrolled on DOL 1, and a provisional diagnosis of Early Infantile Epileptic Encephalopathy type seven was communicated on DOL 4. Carbamazepine was added to levetiracetam(23, 24), providing complete seizure control without sedation, and she was discharged home on DOL 19, in time for a family Christmas. Control neonate 3 was admitted to the same NICU sixteen months earlier, with the same presentation and molecular diagnosis. However, he was diagnosed by standard tests, which took an additional six weeks to return, during which he continued to have uncontrolled seizures. He has severe developmental delay. His 59 day hospitalization cost $211,484. An estimate assuming that the actual cost of $4,426/day for Case 6041 would have persisted for the additional 42 inpatient days, produced estimated rWGS-associated cost savings of $181,481. At eight months of age, she remains seizure free and has minimal developmental delay. The panel consensus was that she avoided likely neurologic damage associated with prolonged uncontrolled seizures(23-27). Cost savings of preventing neurological devastation were not calculated.

Neonate 6053 was transferred to the RCHSD NICU at birth with hypoglycaemia and suggestion that she was the infant of a diabetic mother. She was enrolled on DOL 5, and a provisional diagnosis of focal hyperinsulinemic hypoglycaemia was communicated on DOL 12, indicating the need for surgical evaluation. The median age at surgery for infants with focal hyperinsulinemic hypoglycaemia is 78 days(28). Blood sugars remained brittle during the hospitalization. Persistent or recurrent hypoglycaemia in neonates with hyperinsulinemic hypoglycaemia is associated with neurologic damage, epilepsy, and intellectual disability(9, 28-30). She had surgery on DOL 28, at least 21 days earlier than possible by standard testing(28). The panel consensus was that it was reasonable to compare Case 6053 actual utilization (Figure 1B) vs actual costs plus 21 additional days of NICU stay.

In total, in the six infants, rWGS improved the care of the patients and reduced the costs by at least $803,200 (table 3).The current full cost of rWGS as a CLIA Laboratory Developed Test, including indirect costs and treatment guidance, was $8,482–$22,128 (singleton, duo, trio and quad, table S4). The total cost of rWGS in 42 families was $674,645. Net of rWGS cost in the whole cohort, inpatient cost was estimated to be reduced by $128,555.

## Discussion

The systematic evidence for clinical utility of rWGS in infants with likely genetic diseases, while dramatic, has been limited(9-11). Here, we report the impact of rWGS on clinical outcomes and healthcare utilization in 42 infant inpatients. In contrast to two prior studies, enrolees were predominantly Hispanic/Latino, and enrolled from various inpatient settings, with 71% from intensive care units. 40% received inotropic cardiac support and 76% received ventilatory support. Severity of illness and mortality were similar to the prior study(9). This cohort was well suited to examine clinical utility.

rWGS had a significantly higher diagnostic sensitivity (43%) than standard of care (10%), in agreement with published rates of diagnosis of genetic diseases by WGS (average 40%, range 32% - 57%)(9, 31, 32). Of note, the diagnostic sensitivity was high despite broader enrolment than prior studies(9). This supports the diagnostic sensitivity of rWGS as a first tier test in a substantial subset of infants admitted to children’s hospitals for reasons other than prematurity(31, 33).

The types of genetic diagnoses herein were similar to prior studies of symptomatic infants(7, 9, 34): Autosomal dominant disorders were the most common, and causative variants were de novo as frequently as inherited. There were no recurrent diagnoses, reflecting genetic heterogeneity in inpatient infants.

From blood sample receipt to provisional diagnosis, rWGS is possible in 26–48 hours. Herein, however, median time from enrolment to final report in the medical record was 23 days (range 5–69 days), similar to prior studies(9). Two factors slowed diagnoses: first, confirmatory clinical testing was required before final reporting (adding up to 16 days to time-to-report). Secondly, not all rWGS was performed with the fastest methods, since it was more than two-fold more expensive. In six acutely ill infants (33% of diagnoses), provisional diagnoses were communicated before confirmatory testing (median 7 days, range 3–12 days), since treatments existed that could reduce morbidity or mortality. In the prior two studies, only one infant met these criteria (3% of diagnoses), and diagnosis was in 3 days(9). The clinical utility of rWGS was measured by acute precision medicine interventions, defined as a change in management which physicians agreed resulted directly from the rWGS diagnosis and that occurred promptly after reporting. Exclusive of changes in genetic counselling, subspecialty consultation, or empiric antibiotics, thirteen (72%) of 18 rWGS diagnoses (31% of 42 infants tested) received precision medicine. This concurred with prior studies(9, 33). In a paired test, the rate of clinical utility was significantly greater by rWGS (33%, eleven of 33) than by standard of care (2%, P=0.0015).

The most important measure of clinical utility is improved outcomes. Herein, rWGS-based precision medicine was adjudged to avoid morbidity in 26% (11) of 42 infants, major morbidity in four (10%), and reduced likelihood of acute mortality by 43% in one (2%). Two prior studies reported acute health outcomes of rWGS or exome-based precision medicine in infants. They found that major morbidity was avoided in 3% (four) of 115 infants, and mortality in 3% (three)(9, 33). Two diseases identified in these studies were also seen in our series, with the same avoided morbidity: one was recurrent hypoglycaemia due to focal type 1 familial hyperinsulinism. rWGS enabled expedited partial pancreatectomy, avoiding hypoglycaemic neurologic damage(9, 28-30). The other was Ohtahara syndrome, where rWGS enabled complete control of seizures without sedation, potentially avoiding neurologic damage(9, 23-25, 27). Thus, the magnitude and types of impact of rWGS on outcomes in inpatient infants was replicated in Hispanic/Latino infants in urban southern California.

In addition, the prior study showed rWGS to aid palliative care decisions(9). Herein, diagnosis led to palliative care in one (6%) of 18 infants, less than that study (six (30%) of 20 diagnoses; P=0.09).

When considering introducing new methods in healthcare improved clinical outcomes are the most important factors. However, these outcomes must be considered in the context of health care utilization and cost. To date, studies of genome-wide sequencing have limited financial modelling to compare with the cost of standard genetic tests(34).. Here, we examined healthcare utilization associated with precision medicine interventions in six infants (46% of thirteen receiving rWGS-associated precision medicine). rWGS was estimated to reduce the length of stay by 124 days, and inpatient professional and facility cost by at least $803,200. These estimates are very conservative and reflect the judgement of a Delphi panel. For the palliative care patient the parents elected to end heroic measures once they were aware of the genetic diagnosis. Had this finding occurred earlier in the patient’s care it is possible palliative care would have started earlier saving an additional $1.2 million. This would have made gross savings from the study $2 million. In addition the diagnoses that were made could have very likely been made with proband only analysis which would have yielded a cost of $356,244 for the sequencing.

There were several limitations to the cost analysis. First, it was based on only six “case” infants, two historical “controls”, and published literature values. Second, consensus of a physician panel and the Delphi method were used to judge the impact of precision medicine, and may have been incorrect. Third, utilization was only calculated during that hospitalization, with the exception of a rapid re-admission in infant 6011 and a liver transplant in infant 6026. Children with genetic disorders have substantial ongoing healthcare utilization. We did not include the likely avoidance of a gut transplant in infant 6056, nor avoidance of neurologic damage in infant 6041. Further, infant genetic diseases exert profound emotional, financial, social, and physical stress within families. These include parental divorce, depression and anxiety, and sibling behavioural, developmental, and persistent health complications. The costs of these complications were omitted. Finally, palliative care is not uncommon in regional NICUs, and decisions are often delayed by absence of etiologic diagnoses(7). We did not estimate the optimal rWGS-associated precision medicine scenario, in which rWGS would have been ordered at time of admission.

In conclusion, rWGS is a unique, high-cost healthcare innovation that appears to improve healthcare outcomes while decreasing cost of care. While rWGS merits consideration as a first tier test in a subset of acutely ill inpatient infants, further studies are needed to delineate clinical presentations for which outcomes are consistently improved and optimal timing of rWGS orders and time-to-result.

## Methods

Full methods are described in the Supplementary Material.

### Study Design

Retrospective comparison of clinical utility, outcomes, and healthcare utilization of rWGS and standard of care (including genetic testing) was approved by the institutional review board (IRB) at Rady Children’s Hospital-San Diego (RCHSD) and the Food and Drug Administration (FDA; ClinicalTrials.gov NCT02917460; figure S1). Inpatient infants at RCHSD without etiologic diagnoses, in whom a genetic disorder was possible, were nominated by diverse clinicians from July 26 2016–March 8 2017. Informed consent was obtained from at least one parent or guardian.

### rWGS, Interpretation and Reporting

Clinical features were extracted from electronic medical records (EMR, table S1) and mapped to genetic diagnoses (10, 11). Trio blood samples were obtained where possible. rWGS was performed to ~45-fold coverage as previously described(10, 11). Structural variants were identified with Manta and CNVnator, a combination that provided the highest sensitivity and precision on 21 samples with known structural variants(12, 13). Structural variants were filtered to retain those affecting coding regions of known disease genes and with allele frequencies <2% in the RCIGM database. If WGS established a diagnosis for which a specific treatment was available to prevent morbidity or mortality, this was immediately conveyed to the clinical team. All causative variants were confirmed by Sanger sequencing or chromosomal microarray, as appropriate. Secondary findings were not reported.

### Clinical Utility and Healthcare Utilization

Acute clinical utility of diagnoses and impact on outcomes were evaluated by EMR review, interviews with clinicians, published values, and consensus of at least two paediatricians, one of whom was a relevant paediatric subspecialist and one a medical geneticist. Outcomes were assessed until October 9 2017.

The effect on healthcare utilization was modelled in six infants in whom diagnoses changed treatment and outcomes, by comparing actual healthcare utilization with that of a counterfactual diagnostic trajectory. A modified Delphi method was used to establish consensus for counterfactual trajectories(14). In five infants, the counterfactual trajectory was molecular diagnosis following standard testing based on recent control subjects from the same NICU prior to rWGS and/or literature values. Impact was calculated by the proportionate increase in length of stay multiplied by the actual average daily utilization after one off tests such as MRI scans were excluded. In one infant we considered a counterfactual trajectory as if rWGS was ordered during a prior admission, with correspondingly earlier implementation of precision medicine. In this infant, retrospective impact was calculated by avoidance of the second admission. The cost of rWGS was calculated, including pretest consultation, Sanger confirmation, counselling, result disclosure, and precision medicine guidance (table S4).

### Statistical Analysis

Diagnostic sensitivity and the rate of clinical utility for standard care and rWGS were compared using McNemar’s test for paired nominal data. Fisher’s exact test was used to compare the rate of palliative care initiation with another study. Two-tailed p-values less than 0.05 were considered statistically significant.

## Author contributions

Study concept and design: SFK, DD. Acquisition, analysis or interpretation of data: SC, SN, MC, YD, JAC, MMC, LF, NS, SFK, NV, DD. Drafting of manuscript: SFK, LF, NS, AH, DD. Critical Revision of the Manuscript for Important Intellectual Content: all others. Obtained Funding, Administrative, Technical or Material Support: SFK, JAC. Study Supervision: SFK, JAC, DD, SC.

## Data and material availability

Data are available at LPDR (https://www.nbstrn.org/research-tools/longitudinal-pediatric-data-resource).

## References

1 March of Dimes Foundation Data Book for Policy Makers. Maternal, Infant, and Child Health in the United States 2016. [Available from: http://www.marchofdimes.org/March-of-Dimes-2016-Databook.pdf

2 Xu J, Murphy SL, Kochanek KD, Arias E. Mortality in the United States, 2015. NCHS Data Brief. 2016(267):1–8.

3 Yoon PW, Olney RS, Khoury MJ, Sappenfield WM, Chavez GF, Taylor D. Contribution of birth defects and genetic diseases to pediatric hospitalizations. A population-based study. Arch Pediatr Adolesc Med. 1997;151(11):1096–103.

4 O’Malley M, Hutcheon RG. Genetic disorders and congenital malformations in pediatric long-term care. J Am Med Dir Assoc. 2007;8(5):332–4.

5 Berry MA, Shah PS, Brouillette RT, Hellmann J. Predictors of mortality and length of stay for neonates admitted to children’s hospital neonatal intensive care units. J Perinatol. 2008;28(4):297–302.

6 Petrikin JE, Willig LK, Smith LD, Kingsmore SF. Rapid whole genome sequencing and precision neonatology. Semin Perinatol. 2015;39(8):623–31.

7 Daoud H, Luco SM, Li R, Bareke E, Beaulieu C, Jarinova O, et al. Next-generation sequencing for diagnosis of rare diseases in the neonatal intensive care unit. CMAJ. 2016;188(11):E254–60.

8 Phillips KA, Deverka PA, Sox HC, Khoury MJ, Sandy LG, Ginsburg GS, et al. Making genomic medicine evidence-based and patient-centered: a structured review and landscape analysis of comparative effectiveness research. Genet Med. 2017;19(10):1081–91.

9 Willig LK, Petrikin JE, Smith LD, Saunders CJ, Thiffault I, Miller NA, et al. Whole-genome sequencing for identification of Mendelian disorders in critically ill infants: a retrospective analysis of diagnostic and clinical findings. Lancet Respir Med. 2015;3(5):377–87.

10 Farnaes L, Nahas SA, Chowdhury S, Nelson J, Batalov S, Dimmock DM, et al. Rapid whole-genome sequencing identifies a novel GABRA1 variant associated with West syndrome. Cold Spring Harb Mol Case Stud. 2017;3(5).

11 Hildreth A, Wigby K, Chowdhury S, Nahas S, Barea J, Ordonez P, et al. Rapid whole-genome sequencing identifies a novel homozygous NPC1 variant associated with Niemann-Pick type C1 disease in a 7-week-old male with cholestasis. Cold Spring Harb Mol Case Stud. 2017;3(5).

12 Chen X, Schulz-Trieglaff O, Shaw R, Barnes B, Schlesinger F, Källberg M, et al. Manta: rapid detection of structural variants and indels for germline and cancer sequencing applications. Bioinformatics. 2016;32(8):1220–2.

13 Abyzov A, Urban AE, Snyder M, Gerstein M. CNVnator: an approach to discover, genotype, and characterize typical and atypical CNVs from family and population genome sequencing. Genome Res. 2011;21(6):974–84.

14 Helmer-Hirschberg O. Analysis of the Future: The Delphi Method. Santa Monica, CA: RAND Corporation; 1967.

15 Wymer KM, Anderson BB, Wilkens AA, Gundeti MS. Megacystis microcolon intestinal hypoperistalsis syndrome: Case series and updated review of the literature with an emphasis on urologic management. J Pediatr Surg. 2016;51(9):1565–73.

16 Raphael BP, Nurko S, Jiang H, Hart K, Kamin DS, Jaksic T, et al. Cisapride improves enteral tolerance in pediatric short-bowel syndrome with dysmotility. J Pediatr Gastroenterol Nutr. 2011;52(5):590–4.

17 Lam V, Kain N, Joynt C, van Manen MA. A descriptive report of end-of-life care practices occurring in two neonatal intensive care units. Palliat Med. 2016;30(10):971–8.

18 Oberender F, Tibballs J. Withdrawal of life-support in paediatric intensive care a study of time intervals between discussion, decision and death. BMC Pediatr. 2011;11:39.

19 Lee HP, Kang B, Choi SY, Lee S, Lee SK, Choe YH. Outcome of Alagille Syndrome Patients Who Had Previously Received Kasai Operation during Infancy: A Single Center Study. Pediatr Gastroenterol Hepatol Nutr. 2015;18(3):175–9.

20 Kaye AJ, Rand EB, Munoz PS, Spinner NB, Flake AW, Kamath BM. Effect of Kasai procedure on hepatic outcome in Alagille syndrome. J Pediatr Gastroenterol Nutr. 2010;51(3):319–21.

21 Emerick KM, Rand EB, Goldmuntz E, Krantz ID, Spinner NB, Piccoli DA. Features of Alagille syndrome in 92 patients: frequency and relation to prognosis. Hepatology. 1999;29(3):822–9.

22 Milliman. ’2017 US Organ and Tissue Transplant Cost [Available from: http://www.milliman.com/uploadedFiles/insight/2017/2017-Transplant-Report.pdf.

23 Pisano T, Numis AL, Heavin SB, Weckhuysen S, Angriman M, Suls A, et al. Early and effective treatment of KCNQ2 encephalopathy. Epilepsia. 2015;56(5):685–91.

24 Numis AL, Angriman M, Sullivan JE, Lewis AJ, Striano P, Nabbout R, et al. KCNQ2 encephalopathy: delineation of the electroclinical phenotype and treatment response. Neurology. 2014;82(4):368–70.

25 Kato M, Yamagata T, Kubota M, Arai H, Yamashita S, Nakagawa T, et al. Clinical spectrum of early onset epileptic encephalopathies caused by KCNQ2 mutation. Epilepsia. 2013;54(7):1282–7.

26 Milh M, Boutry-Kryza N, Sutera-Sardo J, Mignot C, Auvin S, Lacoste C, et al. Similar early characteristics but variable neurological outcome of patients with a de novo mutation of KCNQ2. Orphanet J Rare Dis. 2013;8:80.

27 Joshi C, Kolbe DL, Mansilla MA, Mason SO, Smith RJ, Campbell CA. Reducing the Cost of the Diagnostic Odyssey in Early Onset Epileptic Encephalopathies. Biomed Res Int. 2016;2016:6421039.

28 Stanley CA, Thornton PS, Ganguly A, MacMullen C, Underwood P, Bhatia P, et al. Preoperative evaluation of infants with focal or diffuse congenital hyperinsulinism by intravenous acute insulin response tests and selective pancreatic arterial calcium stimulation. J Clin Endocrinol Metab. 2004;89(1):288–96.

29 Hussain K, Blankenstein O, De Lonlay P, Christesen HT. Hyperinsulinaemic hypoglycaemia: biochemical basis and the importance of maintaining normoglycaemia during management. Arch Dis Child. 2007;92(7):568–70.

30 Menni F, de Lonlay P, Sevin C, Touati G, Peigné C, Barbier V, et al. Neurologic outcomes of 90 neonates and infants with persistent hyperinsulinemic hypoglycemia. Pediatrics. 2001;107(3):476–9.

31 Lionel AC, Costain G, Monfared N, Walker S, Reuter MS, Hosseini SM, et al. Improved diagnostic yield compared with targeted gene sequencing panels suggests a role for whole-genome sequencing as a first-tier genetic test. Genet Med. 2017.

32 Stavropoulos DJ, Merico D, Jobling R, Bowdin S, Monfared N, Thiruvahindrapuram B, et al. Whole Genome Sequencing Expands Diagnostic Utility and Improves Clinical Management in Pediatric Medicine. NPJ Genom Med. 2016;1.

33 Stark Z, Tan TY, Chong B, Brett GR, Yap P, Walsh M, et al. A prospective evaluation of whole-exome sequencing as a first-tier molecular test in infants with suspected monogenic disorders. Genet Med. 2016.

34 Stark Z, Schofield D, Alam K, Wilson W, Mupfeki N, Macciocca I, et al. Prospective comparison of the cost-effectiveness of clinical whole-exome sequencing with that of usual care overwhelmingly supports early use and reimbursement. Genet Med. 2017;19(8):867–74.

